# scMultiome analysis identifies embryonic hindbrain progenitors with mixed rhombomere identities

**DOI:** 10.1101/2023.01.27.525932

**Authors:** Yong-Il Kim, Rebecca O’Rourke, Charles G. Sagerström

## Abstract

Rhombomeres serve to position neural progenitors in the embryonic hindbrain, thereby ensuring appropriate neural circuit formation, but the molecular identities of individual rhombomeres and the mechanism whereby they form have not been fully established. Here we apply scMultiome analysis in zebrafish to molecularly resolve all rhombomeres for the first time. We find that rhombomeres become molecularly distinct between 10hpf (end of gastrulation) and 13hpf (early segmentation). While the mature hindbrain consists of alternating odd- versus even-type rhombomeres, our scMultiome analyses do not detect extensive odd versus even characteristics in the early hindbrain. Instead, we find that each rhombomere displays a unique gene expression and chromatin profile. Prior to the appearance of distinct rhombomeres, we detect three hindbrain progenitor clusters (PHPDs) that correlate with the earliest visually observed segments in the hindbrain primordium and that represent prospective rhombomere r2/r3 (possibly including r1), r4 and r5/r6, respectively. We further find that the PHPDs form in response to Fgf and RA morphogens and that individual PHPD cells co-express markers of multiple mature rhombomeres. We propose that the PHPDs contain mixed-identity progenitors and that their subdivision into individual mature rhombomeres requires resolution of mixed transcription and chromatin states.

## INTRODUCTION

Formation of a functioning central nervous system requires that neural progenitors are properly specified and positioned during embryogenesis. One model posits that neural progenitors emerge in segmental compartments (termed neuromeres) that are formed by subdivision of the embryonic neural primordium along its anteroposterior (AP) axis [1]. While it remains under debate whether this model applies to the embryonic fore and midbrain, the embryonic hindbrain is clearly divided into neuromeric units known as rhombomeres [2], but the unique characteristics of each rhombomere have not been defined molecularly and it remains unclear how they form during embryogenesis.

It has long been noted that the hindbrain possesses a two-rhombomere periodicity where each rhombomere pair provides neural crest cells and motor innervation to one specific pharyngeal arch [3, 4]. Each pair consists of one odd- and one even-numbered rhombomere that display unique properties – e.g., neuronal differentiation generally occurs earlier in the even-numbered rhombomere. Transplantation experiments further revealed that cells from odd versus even rhombomeres do not intermix but form discrete boundaries when they come into contact [5]. Strikingly, cells from the even-numbered member of one pair intermix more readily into other even-numbered rhombomeres than into the odd-numbered member of the same pair [6]. Individual rhombomeres are therefore characterized by belonging to a specific rhombomere pair as well as by possessing either an odd or an even identity. This periodicity is recapitulated in some gene expression patterns. For instance, *egfl6* is expressed in even-numbered rhombomeres [7–9] and *ephA4* in odd-numbered ones [10], while *mafba* is expressed in a two-rhombomere unit (r5/r6). r5 and r6 can therefore be assigned to the same rhombomere pair by their shared expression of *mafba* and to an even (*egfl6* expression in r6) versus odd (*ephA4* expression in r5) identity, but similar assignments have not been possible for other rhombomeres. Indeed, relatively few genes have been identified that recapitulate this two-segment periodicity. It is therefore unclear how rhombomeres differ in terms of gene expression programs and whether a two-segment periodicity can be defined molecularly.

It also remains unclear how the rhombomeres emerge from the hindbrain primordium. Visual observations in embryos of various species suggest that rhombomeres develop from broad early segments of the neural tube that are subsequently subdivided. For instance, ‘pro-rhombomeres’ have been observed in the chick [11] and ‘primary’ rhombomeres have been proposed in the human [12, 13]. In the zebrafish, the r3/4 and r4/r5 boundaries form first – thereby delineating r4 – followed by the r6/7 and r1/r2 boundaries that demarcate the r2/r3 and r5/r6 segments. Limited genetic data support the existence of a two-segment pattern in the early hindbrain. Zebrafish *valentino* (*val*) embryos carry a mutation in the *mafba* gene that is expressed in r5 and r6 [14, 15]. In *val* animals, r5 and r6 do not form, but are replaced by an ‘rX’ segment that displays mixed r5/r6 identity. This finding was interpreted to mean that rX represents an early ‘protosegment’ containing progenitors for r5 and r6, but it is unclear if rX cells exist in wildtype embryos and there is no genetic or molecular evidence suggesting the presence of equivalent progenitor domains for other rhombomeres.

Attempts at better defining rhombomere identities and understanding their formation have been hampered by a lack of comprehensive molecular data, as recent scRNAseq analyses could not fully resolve individual rhombomeres [16–20]. We reasoned that scMultiome analysis, which combines RNAseq and ATACseq of individual nuclei, might provide improved definition of rhombomeres. Applying this method in zebrafish, we were able to resolve all rhombomeres for the first time. We find that rhombomeres become molecularly distinct between 10hpf (end of gastrulation) and 13hpf (early segmentation) and remain distinct at 16hpf (mid-segmentation). At these stages, we do not detect extensive similarities among odd versus even rhombomeres, revealing an absence of general molecular identities defining these two states. Prior to the appearance of distinct rhombomeres, our scMultiome analysis identified three cell clusters (HB.1-3) that we refer to as primary hindbrain progenitor domains (PHPDs). Based on our analyses, these PHPDs represent r2/r3 (possibly including r1), r4 and r5/r6. We note that these domains correlate with the earliest visually observed segments in zebrafish embryos and propose that the HB.2 PHPD corresponds to the rX segment identified in *val* mutants. We further find that PHPD cells exhibit mixed gene expression profiles, such that individual cells co-express markers of multiple mature rhombomeres, and we verified the existence of such mixed identity cells in vivo. Lastly, we demonstrate that the retinoic acid (RA) and fibroblast growth factor (Fgf) morphogens are required for formation of the PHPDs. We propose that rhombomeres arise from PHPDs in a process where mixed transcriptional identities are resolved into specific rhombomere fates, but our analyses argue against an early two-segment molecular periodicity and suggest that this pattern may arise later in development.

## RESULTS

### Individual rhombomeres are molecularly resolved at segmentation stages

We applied scMultiome analysis to dissected hindbrain regions at 16hpf (when rhombomeres are becoming morphologically observable; [21]) and at 13hpf (when rhombomere-restricted gene expression is well-established), as well as to whole embryos at 10hpf (the end of gastrulation). After data processing (see Methods), unsupervised clustering followed by projection as UMAPs revealed multiple distinct clusters at each timepoint (Fig. S1-S4). We generated lists of differentially expressed genes for each cluster (Data S2) and, employing a combination of GO term analysis and comparisons to known tissue-specific genes from the literature, we assigned cell type identities to all clusters. Since our samples contain tissues adjacent to the hindbrain primordium, we find clusters corresponding to multiple cell types, but the most prominent group of clusters at each timepoint contained CNS cells. Based on this initial analysis, we bioinformatically isolated CNS-related cell types (excluding placodal cells and post-migratory neural crest cells but including pre-migratory neural crest cells that could not be readily separated), re-clustered them, and projected as UMAPs. These neural UMAPs reveal well dispersed clusters at 13hpf and 16hpf, whereas clusters are more tightly grouped at 10hpf (Figs. 1A-B, 2A-B, 3A-C). In agreement with the early stages being analyzed, most cells in these neural UMAPs express *sox3*, a marker of neural progenitors, with clusters of cells expressing genes indicative of neural differentiation (e.g., *neurod4*) emerging at 13hpf and 16hpf (Fig. 1C-D, 2C-D, 3L-M).

**Figure 1.**
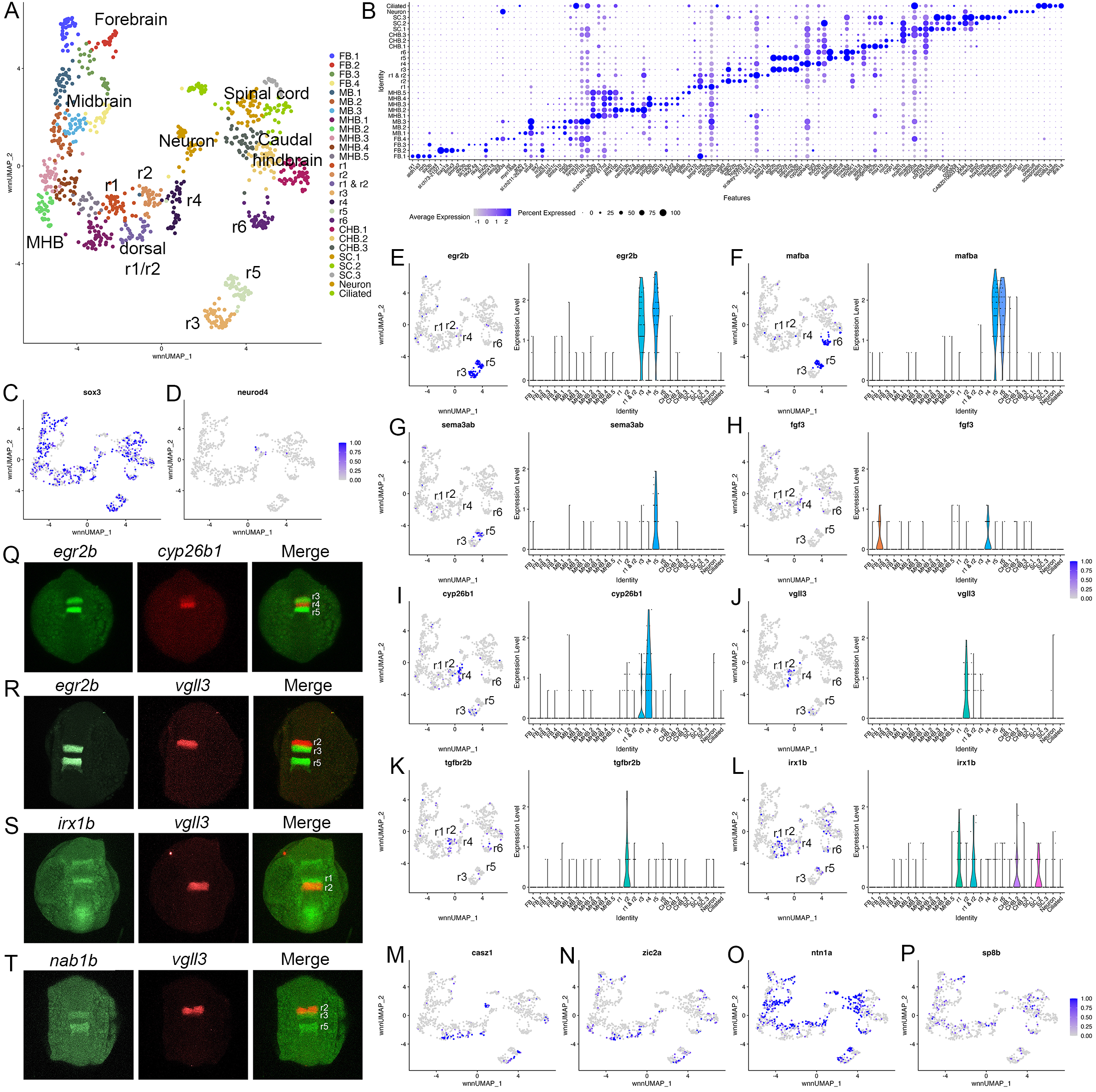
Individual rhombomeres are resolved in the 13hpfzebrafish hindbrain. A. UMAP of 13hpfneural clusters (see text for definition ofneural clusters). B. Dot plot showing expression ofthe top 5 enriched genes in each cluster. C, D. Feature plots showing expressing of *sox3* (C) and *neurod4* (D). E-L. Expression of rhombomere specific genes shown as feature plots (left panels) and violin plots (right panels). M-P. Feature plots showing expression of dorsoventral marker genes. Q-T. HCR analysis of rhombomere-restricted gene expression in 13hpf wildtype zebrafish embryos. Embryos are shown in dorsal view with anterior to the top. FB= forebrain, MB= midbrain, MHB= midbrain-hindbrain boundary, r= rhombomere, CHB= caudal hindbrain, SC= spinal cord.

At early segmentation stages (13hpf), individual rhombomeres are readily identifiable in the neural UMAP based on their gene expression profiles (Fig. 1, Data S2). The r3, r5 and r6 clusters are immediately recognizable based on the known expression of *egr2b* (restricted to r3 and r5; Fig.1E; [22]), *mafba* (restricted to r5 and r6; Fig. 1F; [14]) and *sema3ab* (restricted to r5 at this stage; Fig. 1G; [23]). r4 forms a distinct cluster that is characterized by expression of *fgf3* (enriched in r4; Fig. 1H; [24]), as well as *cyp26b1* that is expressed in r4 before gradually expanding to r3 (Fig. 1I; [25]) – which we confirmed by HCR analysis (Fig. 1Q). A cluster located adjacent to r4 is a strong candidate to contain r2 cells, but there are few known unique markers for r2. Our data reveal that *tgfbr2b* and *vgll3* are highly enriched in this cluster (Fig. 1J, K). A recent report indicated that *vgll3* is expressed in r2 of *Xenopus* [26], but this has not been verified in other systems. We therefore examined *vgll3* expression in 13hpf zebrafish embryos and find that it is restricted to r2, confirming the identity of this cluster (Fig. 1R). A cluster found next to r2 is enriched for *irx1b* (Fig. 1L) and our HCR analysis detected *irx1b* immediately anterior to *vgll3* expression (Fig. 1S), indicating that this cluster corresponds to r1. We note that one remaining cluster at 13hpf expresses markers indicative of both r1 and r2, but this cluster is also enriched in genes broadly expressed in the dorsal neural tube, such as *zic* family TFs (Fig. 1N). A closer examination revealed that dorsal (*casz1* and *zic2a*; [27]) and ventral (*ntn1a* and *sp8b*; [28, 29]) genes are differentially expressed across each rhombomere at 13hpf (Fig. 1M-P), indicating that dorsoventral patterning is ongoing at this stage. Since the dissected hindbrain region used for this analysis includes some adjacent neural tissue, the neural UMAP also contains clusters corresponding to the midbrain-hindbrain boundary (MHB), the caudal hindbrain (CHB), the anterior spinal cord (SC), as well as some cells from the midbrain (MB) and forebrain (FB). An analogous analysis of the 16hpf neural UMAP readily identified r1 to r6 (Fig. 2A, B). At this stage, we observe two clusters that express r5-specific genes (such as *egr2b*). As at 13hpf, we find that dorsoventral markers are differentially expressed between these clusters and across all rhombomeres (Fig. 2E, F).

**Figure 2.**
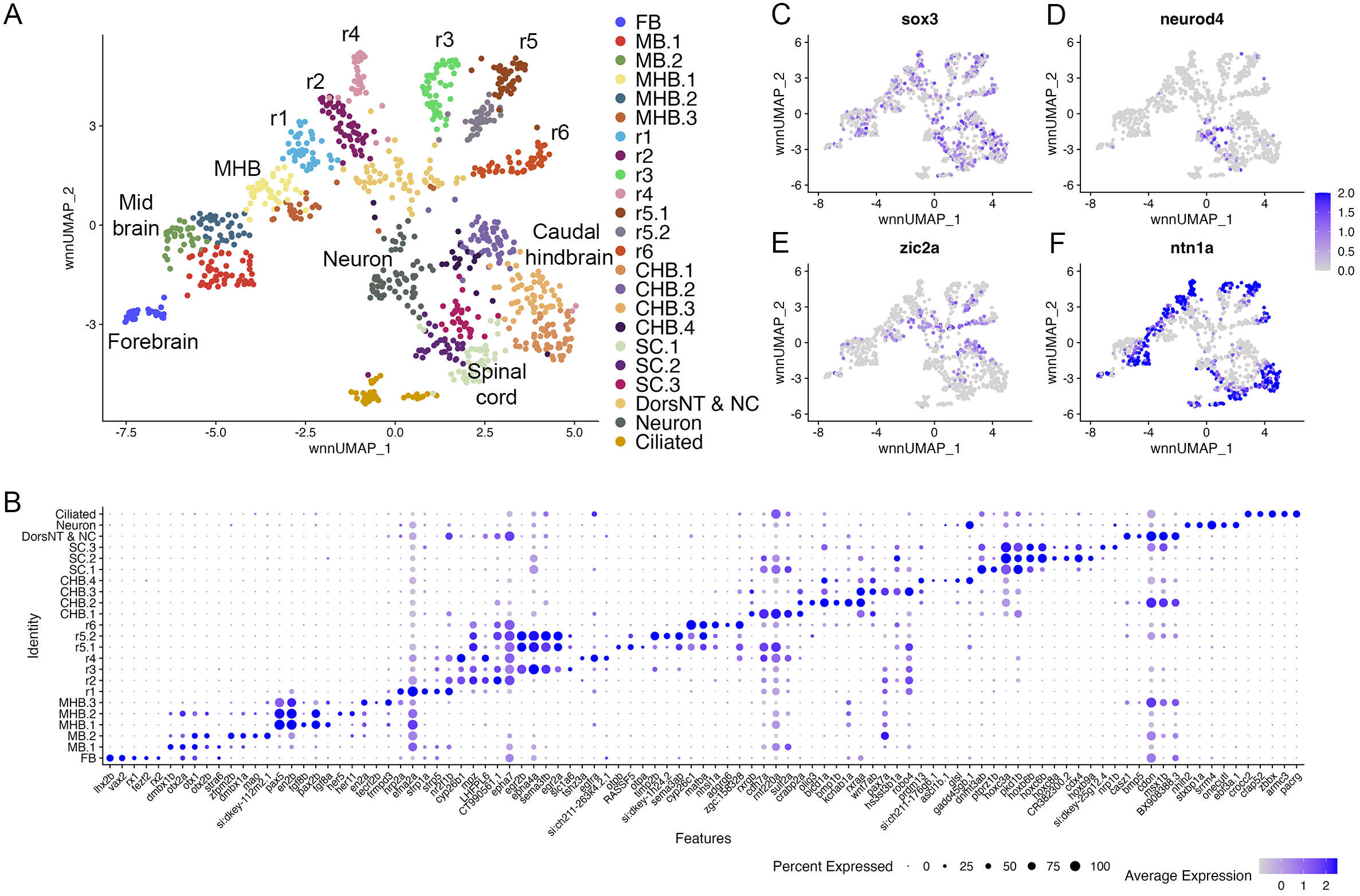
Individual rhombomeres are resolved in the l 6hpf zebrafish hindbrain. A. UMAP of 16hpf neural clusters. B. Dot plot showing expression of the top 5 enriched genes in each cluster. C, D. Feature plots showing expressing of *sox3* (C) and *neurod4* (D). E, F. Feature plots showing expression of dorsoventral marker genes. dorsNT = dorsal neural tube, NC = neural crest. See legend to figure 1 for additional abbreviations.

**Figure 3.**
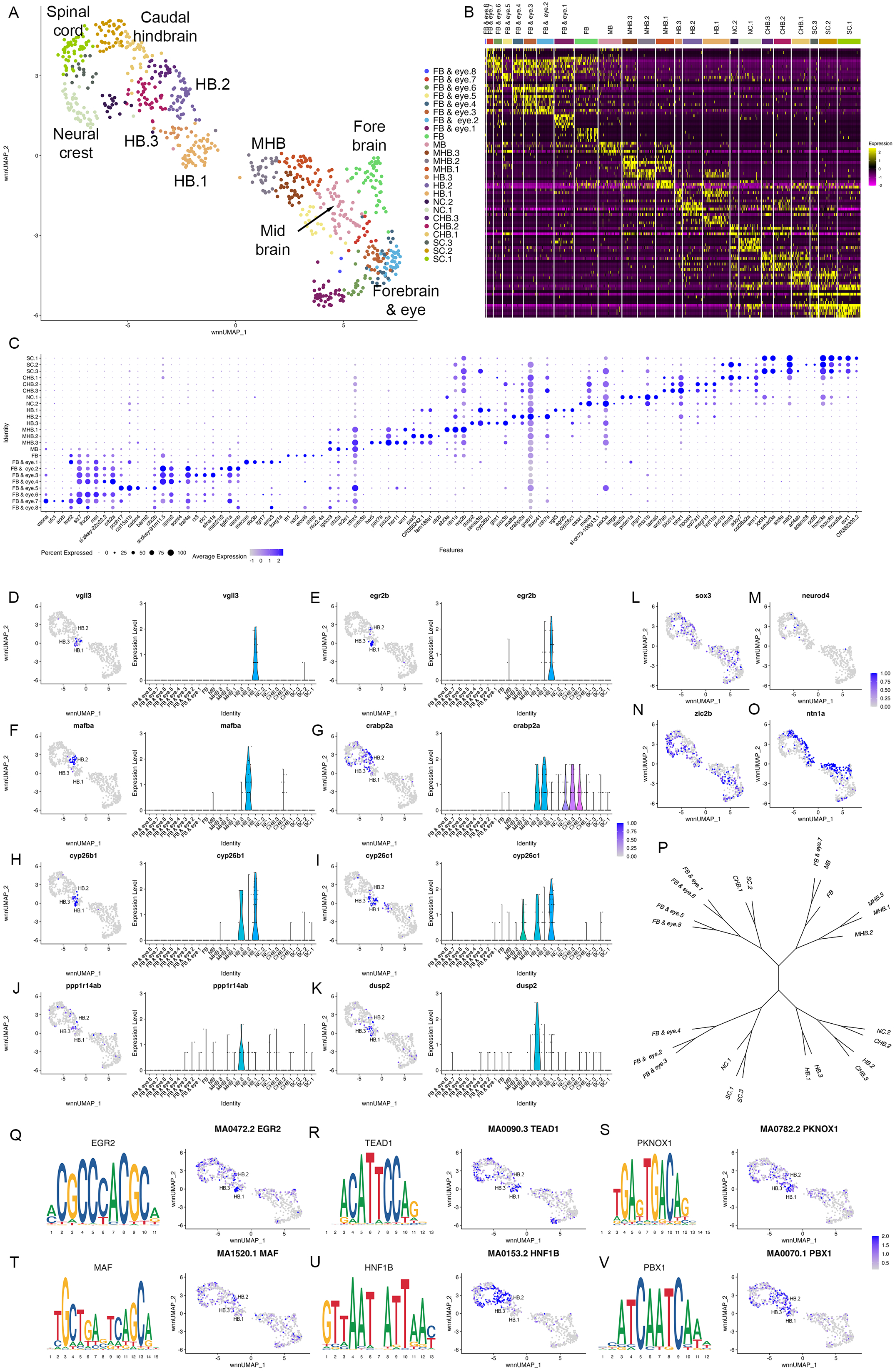
Individual rhombomeres are not resolved at 10hpf in zebrafish. A. UMAP of 10hpf neural clusters. B, C. Heat map (B) and dot plot (C) showing expression of the top 5 enriched genes in each cluster. D-K. Expression of rhombomere-specific genes shown as feature plots (left panels) and violin plots (right panels). L-M. Feature plots showing expressing of *sox3* (L) and *neurod4* (M). N, O. Feature plots showing expression of dorsoventral marker genes. P. Dendrogram showing the relationship between the 10hpf neural clusters. Q-V. Rhombomere-enriched accessible transcription factor binding motifs are shown as a motif logo (left panels) and as a feature plot of chromVar activity (right panel). HB = hindbrain, NC = neural crest. See legend to figure 1 for additional abbreviations.

To further confirm the assigned rhombomere identities, we identified ATAC peaks enriched in each cell cluster relative to all other clusters and carried out a de novo search for over-represented transcription factor (TF) binding motifs in these differentially accessible regions (Data S3, Fig. 4). At both 13hpf and 16hpf, the r3 and r5 clusters display a strong enrichment for accessible EGR2 motifs (Fig. 4C; consistent with high expression of *egr2b* in both rhombomeres), but the r3 and r5 clusters also differ from each other such that the r5 cluster is enriched for accessible MAF motifs (Fig. 4D; in agreement with *mafba* expression in r5) and r3 is enriched in accessible binding motifs for TALE TFs (such as PKNOX1 [a.k.a., Prep1]; Fig. 4E). The type of TALE binding motif observed in r3 is also enriched in other anterior rhombomeres, and corresponds to a compound binding site that supports binding of TALE heterodimers (Pbx:Prep and/or Pbx:Meis; [30, 31]), which are broadly expressed in the hindbrain [32–37]. r6 shows enrichment for accessible MAF motifs (Fig. 4D; in accordance with *mafba* expression in r6), as well as for paralog group (PG) 4 HOX motifs (Fig. 4F). PG4 HOX motifs are also enriched in the caudal hindbrain at 13hpf and become less accessible at 16hpf, consistent with the dynamic expression of several PG4 *hox* genes in the caudal CNS with anterior limits at r5 and r6 [38, 39]. The r4 cluster is enriched for Pbx binding motifs that are also accessible in some more caudal tissues (Fig. 4G). Notably, this type of Pbx site is distinct from the heterodimeric TALE sites enriched in r3 and are reported at enhancers where Pbx TFs act as co-factors to Hox proteins – such as at several Hox-dependent enhancers active in r4 [40–45]. Cells in the r2 cluster are highly enriched for accessible binding motifs for TEAD family TFs at 13hpf (Fig. 4H). TEAD TF expression is not restricted to r2 (Fig. 4A, B), but TEAD TFs are known heterodimerization partners to Vgll TFs [46, 47] and *vgll3* expression is restricted to r2 (Fig. 1J, R-T). This suggests that Vgll3 supports binding of TEAD TFs to genomic DNA specifically in r2. We did not identify any TF binding motifs unique to r1, but the r1 and r2 clusters (along with some MHB clusters) display enrichment for NR2C/F family motifs (Fig. 4I, J), consistent with *nr2f1b* being expressed in this region of the zebrafish embryo [48]. Lastly, we note that many motifs enriched in the rhombomeres are also enriched in clusters containing differentiating neurons, suggesting that some genomic regions remain accessible as progenitors begin to differentiate. Overall, we find strong agreement between the scRNA and scATAC data in defining individual rhombomeres and we conclude that r1-r6 are molecularly distinct from each other and adjacent neural structures by 13hpf, which precedes the stage when individual rhombomeres become visually detectable in zebrafish.

**Figure 4.**
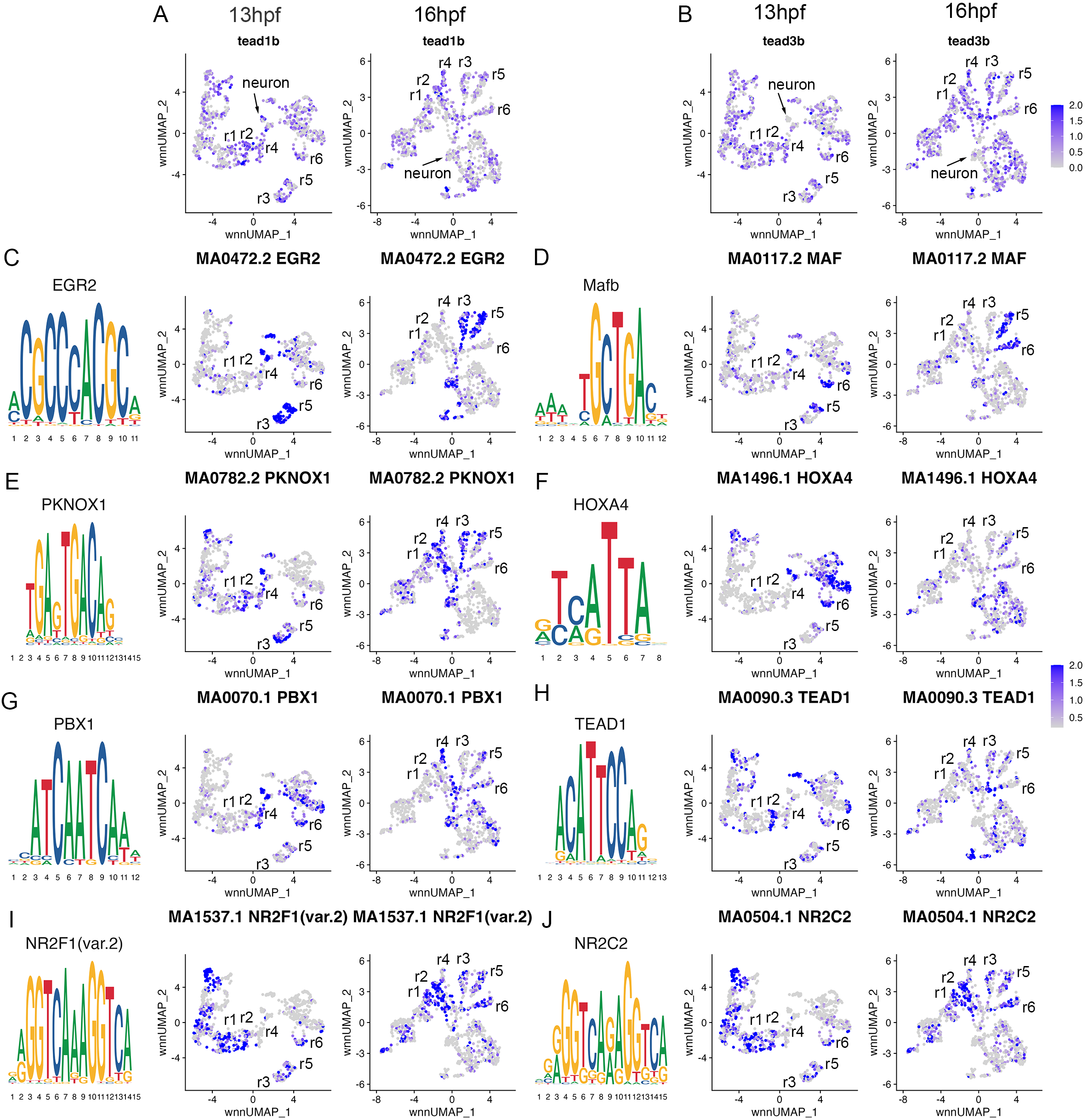
Transcription factor binding motifs show rhombomere-restricted accessibility. A, B. Feature plots showing expression of TEAD transcription factors at 13hpf and 16hpf. C-J. Rhombomere-enriched accessible transcription factor motifs are shown as a motif logo(left panels) and as a feature plot of chromVar activity at 13hpf(middle panel), or 16hpf(right panel).

### The early hindbrain does not display a molecular rhombomere periodicity

The mature hindbrain appears to possess a periodicity, such that pairs of one odd- and one even-numbered rhombomeres are arranged along its anterior-posterior axis. This concept is based on relatively mature characteristics of the hindbrain, such as the exit of branchiomotor nerves, the migration of neural crest cells and the onset of neuronal differentiation [3, 4]. Despite the discovery of some genes that recapitulate this pattern (e.g., *ephA4a* expression in r1/r3/r5 [10], and *egfl6* expression in r2/r4/r6 [8]), it is unclear if this periodicity is present earlier in development and if it is broadly recapitulated molecularly – in the form of gene expression and chromatin accessibility patterns. The UMAPs presented in Figs. 1 and 2 do not show odd and even rhombomeres clustering into separate groups. Instead, r1, r2 and r4 cluster with anterior neural structures, while r6 clusters closer to caudal structures. Since the position of clusters in the UMAP are not fully indicative of how related two cell populations are, we next generated heatmaps of the top genes in each cluster (Fig. 5A, B). The heatmaps reveal considerable gene expression overlap between r3 and r5, but these rhombomeres do not share extensive expression with r1. Similarly, while r2, r4 and r6 share some gene expression, r2 seems to express more genes related to r3, and r6 shares closer expression with r5. To examine these relationships more comprehensively across all rhombomeres, we used the weighted nearest neighbor graphs from the Seurat analysis and projected them as dendrograms (Fig. 5C, D). This analysis revealed very similar relationships at 13hpf and 16hpf. At both stages, r3 and r5 are closely related to each other, while r1 is more closely related to the MHB (13hpf) or r2 (16hpf). Further, r4 is related to r2, but not to r6, which instead clusters with the caudal hindbrain at both stages. As a result, there is no clustering of even versus odd rhombomeres at either 13hpf or 16hpf. To more directly assess if there is a molecular odd versus even pattern among rhombomeres, we computationally identified any genes expressed in r2, r4 and r6 – and whose average expression in those rhombomeres exceeds their expression in r1, r3 and r5 – or vice versa at 13hpf (Fig. 5E, F). We find very few genes that fulfill these criteria. *col7a1l* is highly expressed in r2, r4 and r6, but most other genes expressed in an even or odd pattern are expressed weakly in one of the rhombomeres. For instance, *plxna2* is robustly expressed in r2 and r4, but only weakly in r6, while *egr2b, ephA4a* and *sema3fb* are highly expressed in r3 and r5, but barely detectable in r1. We carried out an analogous analysis to identify genes expressed in two adjacent rhombomeres, to determine if rhombomere pairs can be defined by co-expression of genes. This analysis identified several genes co-expressed in r5 and r6 (Fig. 5I), like the known expression of *mafba* [14] and the PG3, 4 *hox* genes [38, 39]. In contrast, we detect only a few genes co-expressed in r1 and r2 (*nrp2a* and *pdzrn4*; Fig. 5G) and none co-expressed in r3 and r4. We conclude that there is not a strong molecular odd versus even, or rhombomere pair, molecular pattern (except for the r5/r6 pair) during rhombomere formation, indicating that the odd versus even features observed in the mature hindbrain may instead be achieved by later-acting mechanisms.

**Figure 5.**
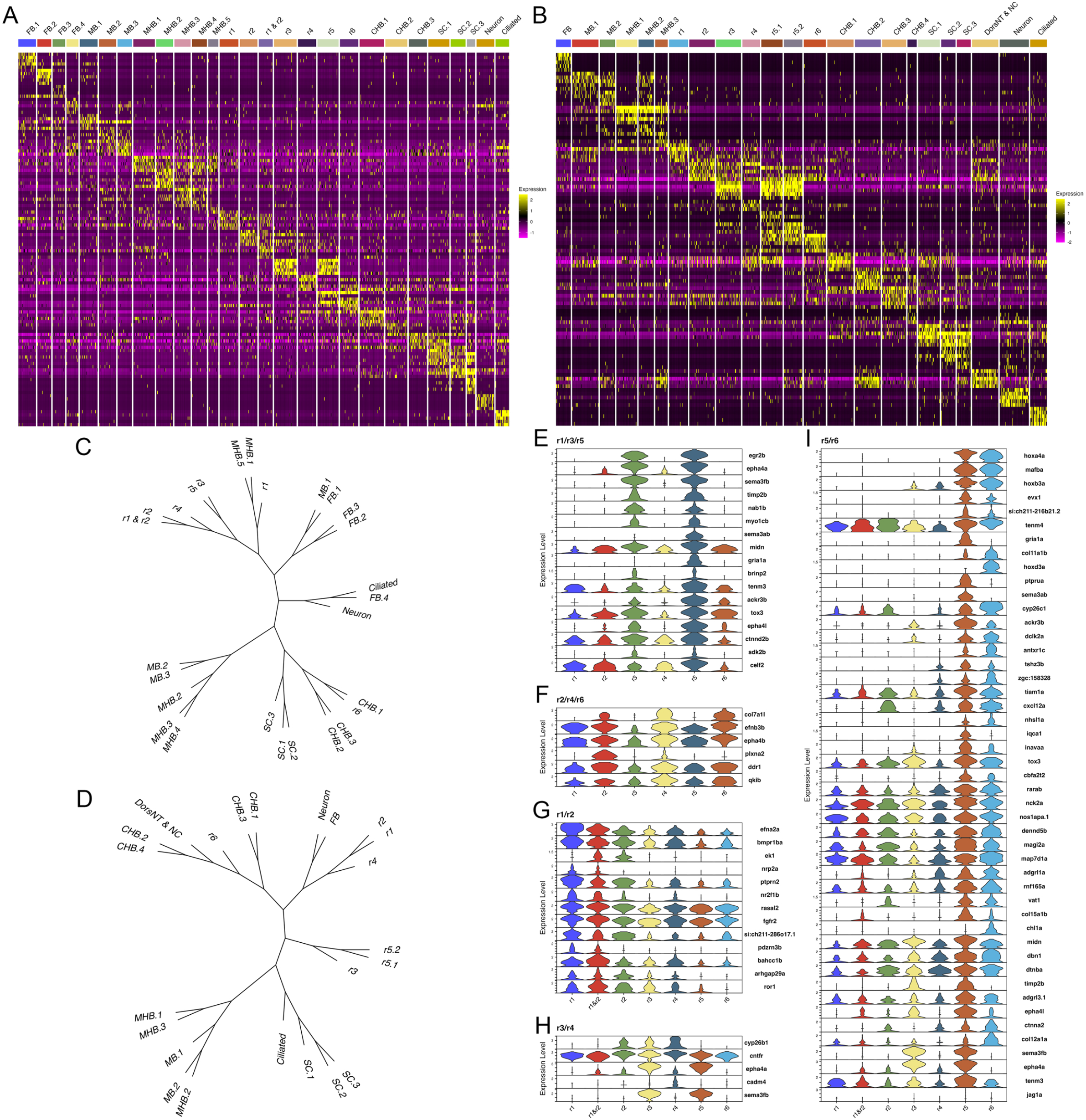
The early hindbrain lacks a detectable odd versus even rhombomere pattern. A, B Heat maps showing expression of the top 5 enriched genes in each cluster at 13hpf (A) and 16hpf (B). C, D. Dendrograms showing the relationship between each neural cluster at 13hpf (C) and 16hpf (D). E-I. Violin plots showing expression levels of genes enriched in odd (E) or even (F) rhombomeres, as well as genes enriched in adjacent pairs of rhombomeres (G-I), at 13hpf.

### scMultiome identifies three hindbrain progenitor domains at the end of gastrulation

To determine when rhombomeres emerge during embryogenesis, we examined the 10hpf neural UMAP. Using gene expression (Data S2) and motif accessibility (Data S3), we assigned cell identities to each cluster (Fig. 3A-C). We find that, in contrast to 13hpf and 16hpf, individual rhombomeres are not apparent as distinct clusters at 10hpf. Instead, we identified three clusters (HB.1-HB.3) that express genes characteristic of mature rhombomeres. HB.1 is the largest of these and expresses genes associated with several different rhombomeres such that *vgll3* (Fig. 3D; an r2 marker; Fig. 1R; [26]) and *egr2b* (Fig. 3E; a marker of r3 and r5; [22]) are both enriched in this cluster. A second prominent rhombomere cluster (HB.2) expresses *mafba* (Fig. 3F; a marker of r5/r6; [14]) and *crabp2a* (Fig. 3G; expressed in r6, the caudal hindbrain, and weakly in r4/r5; [49]), as well as limited *egr2b* (Fig. 3E). Since *egr2b* expression is initiated in r3 at the end of gastrulation (10hpf), but does not become expressed in r5 until 11hpf, this suggests that HB.1 and HB.2 correspond to prospective anterior and posterior rhombomeres, respectively. Furthermore, a third rhombomere cluster (HB.3) is related to HB.1 as these two clusters co-express *cyp26b1* and *cyp26c1* (Fig. 3H, I; expressed in r2-r4; [25, 50]), but HB.3 does not express either *vgll3* or *egr2b* (Fig. 3D, E). Instead, HB.3 expresses *dusp2* and *ppp1r14ab*, two markers that we previously reported as restricted to r4 at this stage (Fig. 3J, K; [8, 51]). All HB clusters are clearly separate from a group of clusters corresponding to the MHB, the midbrain and the forebrain, while the HB.2 cluster is adjacent to several clusters corresponding to the caudal hindbrain and the anterior spinal cord. TF motif accessibility generally supports the gene expression analysis, and we find that multiple TF binding motifs are associated with the HB clusters at 10hpf. Egr, TEAD and TALE binding motifs are enriched in accessible chromatin regions in HB.1 cluster cells (Fig. 3O-Q), which is supportive of our assertion that HB.1 contains precursors of anterior rhombomeres. Similarly, the HB.2 cluster is enriched for accessible Maf and Hnf1 motifs in accordance with it containing posterior progenitors (Fig. 3R, S). Egr motifs are also weakly enriched in HB.2, consistent with the *egr2b* gene becoming activated in r5 at this stage. Lastly, we find that Pbx motifs are enriched in HB.3 (Fig. 3T). Although Pbx motifs are also enriched in HB.1 and HB.2, HB.3 is unique in that it shows minimal enrichment for accessible Egr, TEAD, Hnf1 and Maf motifs. These relationships are supported by dendrogram projections (Fig. 3N), where HB.2 clusters with the caudal hindbrain, whereas HB.1 and HB.3 cluster together on a separate branch. Taken together, this analysis demonstrates that individual rhombomeres are not yet molecularly distinct at 10hpf. Instead, three progenitor domains with different anteroposterior characteristics are present, such that HB.1 likely corresponds to the future r2/r3 (and possibly r1), HB.2 to r5/r6 and HB.3 to r4. We tentatively refer to these domains as primary hindbrain progenitor domains (PHPDs).

### The PHPDs contain progenitors with mixed rhombomere identities

To determine how the PHPDs relate to more mature rhombomeres, we integrated neural clusters from all three timepoints. To simplify this analysis, we omitted all forebrain cells and projected the remaining cells as a UMAP. The integrated UMAP reveals clearly delineated clusters, and we identified each rhombomere based on its gene expression and TF motif accessibility profile (Data S4, S5), as we did for the individual timepoints (Fig. 6A). We again find that each rhombomere is split into dorsal (expressing *zic2b*) and ventral (marked by *ntn1a*) clusters – except for r3, which is represented by a single cluster that nevertheless shows differential expression of dorsoventral genes in distinct subdomains (Fig. 6B, C). Projecting the integrated data in the form of a dendrogram (Fig. 6D) largely confirms the relationships observed at individual timepoints such that r1 and r2 cluster together with r4 on a shared branch, r3 and r5 cluster on a separate branch, and r6 forms its own branch. We then used the integrated UMAP to examine the contribution of cells from each timepoint to the integrated rhombomere clusters (Fig. 6E; Data S6). We find that cells from individual rhombomeres at 13hpf and 16hpf combine into the integrated clusters such that, for instance, r3 cells from 13hpf and 16hpf contribute to the integrated r3 cluster (Fig. 6E), indicating that hindbrain cells have shared rhombomere characteristics at 13hpf and 16hpf. In contrast, cells from the 10hpf PHPD clusters do not contribute fully to the integrated rhombomeres. A portion of cells from the 10hpf HB.2 cluster distribute to the integrated r5 and r6 clusters, but many remain as a separate cluster (HB) in the integrated UMAP (Fig. 6E), indicating that – although they express *mafba* (a marker of r5/r6; Fig 3F) – their overall molecular profile is sufficiently distinct from more mature r5/r6 cells that they remain as a separate group. Even the HB.2 cells that do distribute to rhombomeres are actually located between r4 and r6 in the integrated UMAP, again indicating that they are distinct from more mature r6 cells. Similarly, while cells from the 10hpf HB.1 cluster distribute to the integrated r1, r2 and r3 clusters (Fig. 6E), a detailed examination of their distribution reveals that many occupy a position between r1 and r2, further indicating that PHPD cells are molecularly distinct from more mature rhombomere cells.

**Figure 6.**
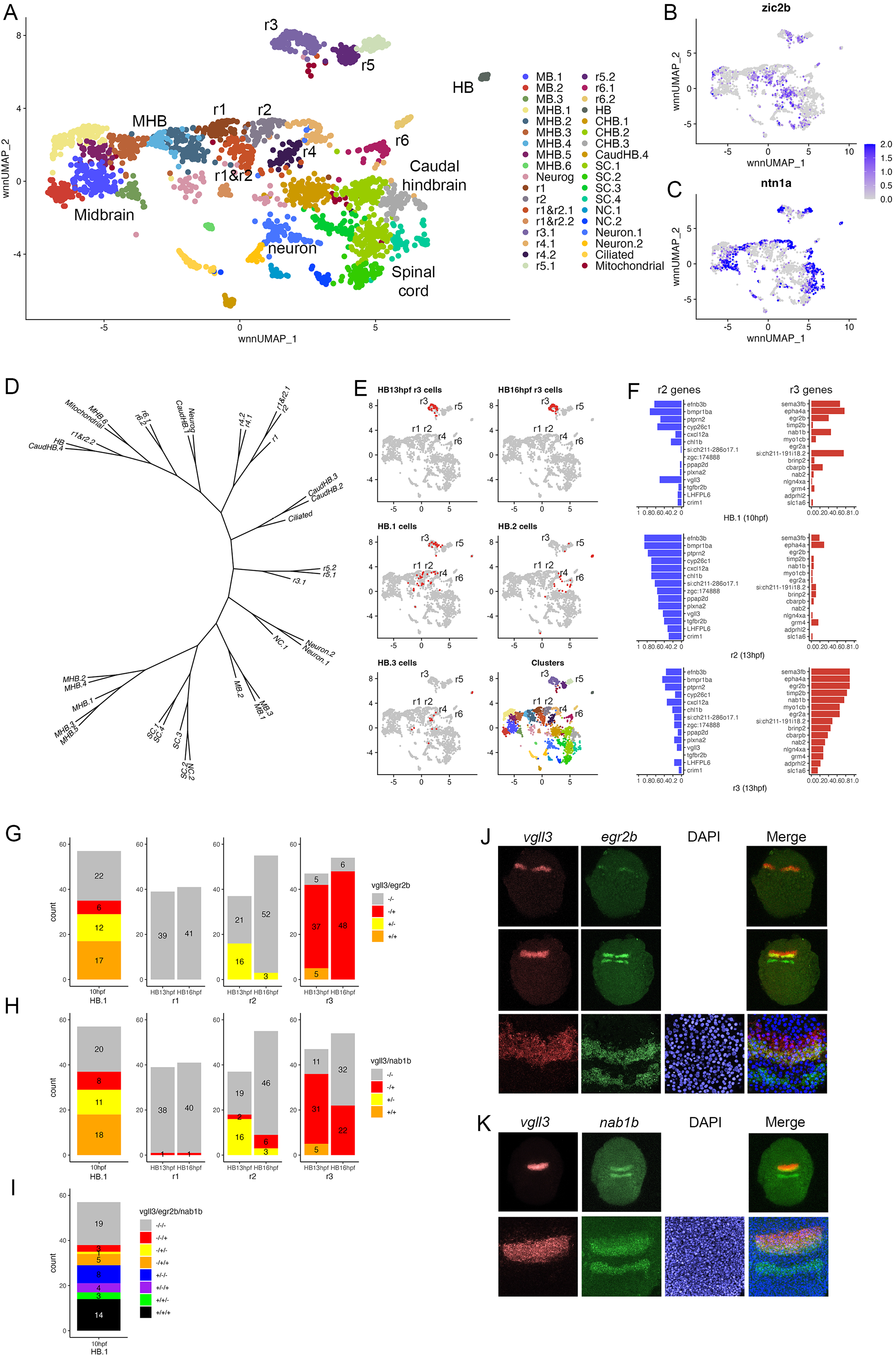
Rhombomere progenitor cells display mixed gene expression identities. A. UMAP of integrated data from 10hpf, 13hpf and 16hpf. B, C. Feature plots showing expression of dorsoventral marker genes. D Dendrogram showing the relationship between the integrated neural clusters. E. Feature plots showing contribution of various cell populations (red cells) to the integrated rhombomere clusters. F. Bar graphs showing expression of r2 genes (left column) and r3 genes (right column) in 10hpf HB.1 cells (top panel), 13hpf r2 cells (middle panel) and 13hpf r3 cells (bottom panel). G-I. Co-expression of r2 markers (*vgll3*) and r3 markers (*egr2b, nab1b*) in individual cells from HB.1, r1, r2 and r3. Numbers in the bar graphs represent cell counts. J-K. HCR analysis of rhombomere-restricted gene expression in 10-11hpf wildtype zebrafish embryos. Embryos are shown in dorsal view with anterior to the top. The bottom row in each panel displays a higher magnification of the row above.

To better characterize the transcriptional state of PHPD cells, we examined the HB.1 cluster more closely. We identified the top genes differentially expressed in r2 versus r3 at 13hpf (and vice versa). As expected, these genes are preferentially expressed in their corresponding rhombomere at 13hpf (Fig. 6F, middle and bottom panels), but we find that both r2 and r3 genes are extensively expressed in HB.1 cells at 10hpf (Fig. 6F, top panel). Examining the expression state of individual HB.1 cells, we identified many cells that express both *vgll3* (r2 marker) and *egr2b* (r3 marker at this stage), such that ~74% of *egr2b*-expressing cells co-express *vgll3* at 10hpf (Fig. 6G; left panel). This contrasts with 13hpf – when r2 cells do not co-express *egr2b* and only ~10% of r3 cells co-express *vgll3* – and 16hpf, when *egr2b* is fully restricted to r3 and *vgll3* is being downregulated. In addition to co-expressing *egr2b*, we find that *vgll3-*positive cells also co-express *nab1b* (r3 and r5 marker; [52]; Fig. 1T) such that 62% of *vgll3* positive cells co-express *nab1b* (Fig. 6H). When considered in combination, ~72% of *vgll3-*positive cells co-express either *egr2b* or *nab1b* at 10hpf (Fig. 6I). To identify the HB.1 domain and assess co-expression of rhombomere markers in vivo, we turned to HCR analysis in zebrafish embryos. We find extensive overlap between *vgll3* and both *egr2b* and *nab1b* expression at the earliest stages that these genes can be detected (Fig. 6J, K). The PHPDs therefore differ from later rhombomeres in terms of their molecular profile and, at least in the case of HB.1, individual PHPD cells possess mixed identities characterized by co-expression of markers for multiple rhombomeres.

### Morphogens control formation of the PHPDs

The early hindbrain primordium is exposed to morphogens in vivo, with RA acting posteriorly (from its source in the adjacent somites) and Fgfs acting anteriorly (with an initial source at the MHB followed by a secondary source in r4), to establish an anteroposterior pattern (reviewed in [53, 54]). Previous work has demonstrated that various degrees of RA signal disruption produces distinct effects on hindbrain gene expression (reviewed in [55]), but complete loss of RA signaling appears to disrupt formation of the caudal rhombomeres [56], suggesting that RA may be required for formation of the HB.2 domain. Accordingly, we find that expression of RA signaling components – such as the *RARab* retinoic acid receptor and the *crabp2a* retinoic acid binding protein – is enriched in HB.2 cells (Fig. 7A, B). To test if formation of the HB.2 domain requires RA signaling, we used DEAB (an inhibitor of RA synthesis) to block RA signaling and assessed early HB.2 gene expression by HCR. We find that *egr2b* expression in r5 and *mafba* expression in r5/r6 first becomes detectable by 10hpf-11hpf in control embryos, but these expression domains are never observed in DEAB-treated embryos (Fig. 7H-K). In contrast, expression of *vgll3*, *cyp26b1* and *cyp26c1*, as well as the anterior *egr2b* expression domain, persists in DEAB-treated embryos, but expression of these genes is observed over a broader region and may also be somewhat weaker (Fig. 7H-K). Previous work demonstrated that disruption of r5/r6 formation in *mafB* mutant embryos results in a unique rX cell population that is positioned between r4 and the caudal hindbrain [14, 15]. To determine if such an rX domain forms in place of r5/r6 in DEAB-treated embryos, or if this region may be lost, we assessed the expression of *cdx4* in the spinal cord relative to *cyp26b1* in r4 and *egr2b* in r3 (Fig. 7L, M). As in Fig. 7H-K, we again observe a slight expansion of *egr2b* in r3 and *cyp26b1a* in r4 of DEAB-treated embryos. These anterior expression domains closely abut *cdx4* expression without an apparent rX region. In agreement with a role also for Fgf signaling, we detect expression of Fgf8 and Fgf3 in several PHPD clusters of the 10hpf UMAP (Fig. 7C, D), and we find that treatment with SU5402 (an inhibitor of Fgf receptor signaling) blocks expression of *vgll3* and *egr2b* in r2/r3 at 10hpf (Fig. 7N) – as well as of *mafba* and *egr2b* in r5/r6 and of *hoxb1a* in r4 – when those genes first become detectable at 12hpf-13hpf (Fig. O, P). Therefore, while RA signaling is specifically required for HB.2 formation, Fgf signaling appears broadly required to establish the PHPDs.

**Figure 7.**
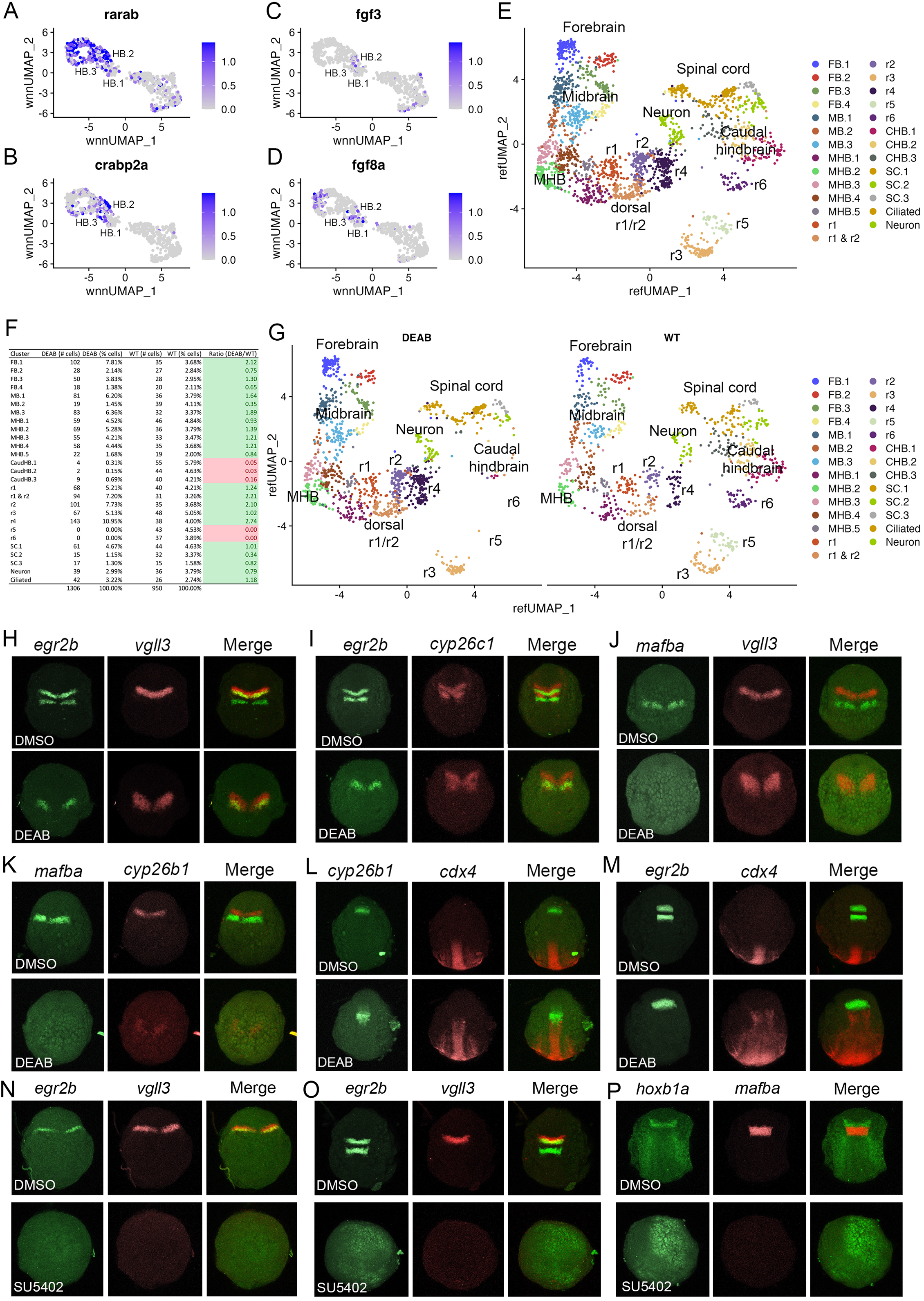
PHPD formation is controlled by morphogens. A-D. Feature plots showing expression of RA and Fgf signaling components at 10hpf. E. Combined UMAP of 13hpf wildtype and DEAB-treated embryos. F. Contribution of wildtype (WT) and DEAB-treated cells to each cluster in E. G. Separate UMAP projections of wildtype (right panel) and DEAB-treated (left panel) embryos at 13hpf. H-M. HCR analysis of rhombomere-restricted gene expression in control (top panels) and DEAB-treated (bottom panels) embryos at 10-11hpf. N-P. HCR analysis of rhombomere-restricted gene expression in control (top panels) and SU5402-treated (bottom panels) embryos at 10-11hpf (N) or 12-13hpf (O-P). Marker gene expression: *egr2b*: r3/5; *vgll3*: r2; *cyp26c1*: r2/4; *mafba*: r5/r6; *cyp26b1*: r4; *hoxb1a*: r4; *cdx4*: spinal cord. Embryos are shown in dorsal view with anterior to the top.

While these analyses confirm a role for morphogens in inducing hindbrain gene expression and suggest that RA and Fgf may be required for formation of the PHPDs, they rely on HCR analyses of a small number of genes. It is therefore possible that expression of other rhombomere markers persists and/or that the remaining rhombomeres are mis-specified in embryos with disrupted morphogen signals. Previous analyses of morphogen function in early hindbrain development suffer from the same imitations, as they also did not comprehensively assess effects on rhombomere gene expression. To address this shortcoming, we again turned to scMultiome analysis. Since RA signaling appears to be specifically required for HB.2 formation we focused on this morphogen, collected DEAB-treated embryos at 13hpf (Fig. S3; Data S2), and combined DEAB-treated with control samples into an integrated UMAP (Fig. 7E). In assessing the contribution of control versus DEAB-treated cells to each UMAP cluster, we find that the r5 and r6 clusters contain only cells from control embryos and no cells from DEAB-treated embryos (Fig. 7F). Accordingly, projecting DEAB-treated and wild type samples in separate UMAPs reveals a complete loss of r5 and r6 in the DEAB condition (Fig. 7G), consistent with the HB.2 progenitor population not forming. In agreement with the apparent expansion of anterior rhombomeres observed in our HCR analyses of DEAB-treated embryos, cell ratio comparisons reveal that r1 and r2 contain twice as many cells, and r4 contains almost three times as many cells, in DEAB-treated compared to control embryos (Fig. 7F). Furthermore, the caudal hindbrain (CHB) clusters are lost in DEAB-treated embryos while the spinal cord clusters largely persist (Fig. 7F, G). We also find that all cells from DEAB-treated embryos can be mapped to a wildtype cluster, again indicating that an rX domain does not form in DEAB-treated embryos. Further, we find only minor changes in gene expression in r1-r4 upon loss of RA signaling (Fig. S5; Data S7). Only ~30 genes are differentially expressed between DEAB-treated and control embryos (>2-fold change; pval<1e-5) in each rhombomere – and many of these genes are shared between rhombomeres – indicating that they reflect broadly RA-responsive genes rather than genes that affect individual rhombomere fates. Our results suggest that RA is required for formation of the HB.2, but not the HB.1 or HB.3 domains, although we cannot exclude the possibility that a small number of genes are RA-dependent in the anterior domains.

## DISCUSSION

Hindbrain rhombomeres represent a well-studied model for neural progenitor specification and positioning, but we still lack a clear understanding of the molecular basis for differences between rhombomeres, and we do not know how they form in the embryo. This is largely due to a lack of comprehensive molecular data as individual rhombomeres have not been resolved by previous scRNAseq analyses on whole embryos [16–18, 20], or on dissected hindbrain regions [19]. Using combined single nucleus RNAseq and ATACseq (scMultiome), we successfully resolved all rhombomeres at zebrafish segmentation stages. We identify a unique molecular profile for each rhombomere and find that there is no clear two-segment periodicity to rhombomere identities at these stages. We also define three mixed-identity progenitor domains that likely correspond to the pre-rhombomeres observed visually in early embryos of several species and that are thought to subsequently subdivide into the mature rhombomeres. Our results therefore unify previous visual observations with a comprehensive molecular characterization into a general mechanism for rhombomere formation.

### The early rhombomeres lack an overt molecular periodicity

The maturing hindbrain possesses a two-segment periodicity [3, 4], but the molecular basis of this pattern is not clear, and it is not known how early it may be present. Using the full gene expression profile for each rhombomere at segmentation stages, we find that r3 and r5 are closely related and that r2 shares features with r4, but r1 and r6 do not fall into these groups (Fig. 5A-D). Indeed, we find very few genes that are restricted to odd versus even rhombomeres at these stages (Fig. 5E, F). Similarly, while we observe numerous genes co-expressed in r5 and r6, we do not detect shared gene expression in other pairwise combinations of rhombomeres (Fig. 5G-I). These results indicate that repeating two-segment molecular identities are not overtly present in the early zebrafish hindbrain.

Instead of a molecular periodicity, each rhombomere displays a distinct molecular profile and its relatedness to other rhombomeres changes over time. For instance, r3 and r5 progenitors reside in the HB.1 and HB.2 domains, respectively, at 10hpf and these populations do not show extensive overlap in gene expression, but r3 and r5 cells are closely related by 13hpf. This means that r3 and r5 progenitors must diverge from their closest neighbors at 10hpf (r2 and r6 progenitors, respectively) and converge on a shared gene expression program by 13hpf. This process is likely driven by the action of TFs shared between r3 and r5. Accordingly, *egr2b* is one of the earliest TFs expressed in r3 and r5, and separate regulatory elements required for its expression in each rhombomere have been identified [57–62]. A similar effect may be mediated by *mafba* in r5 and r6 [14, 63], as these rhombomeres co-express several genes. Notably, r5 clusters with r3, not r6, in the dendrograms suggesting that *egr2b* may exert a stronger effect than *mafba*, perhaps because these TFs control gene networks of different size and/or composition.

Although the early rhombomeres do not display an odd versus even molecular periodicity, they nevertheless possess some odd versus even features. Transplantation and cell mixing experiments have revealed that cells from odd rhombomeres mix more readily into other odd rhombomere than into even ones, and vice versa [5] – a process likely controlled by cell surface molecules such as the *ephrins* and their *Eph* receptors [10, 64, 65]. This finding raises the possibility that expression of cell surface receptors may be controlled by gene networks shared between odd versus even rhombomeres. If such networks are small, they might be operating without giving rise to detectable molecular signatures in our analyses. However, previous examination also revealed a hierarchy for cell mixing such that, for instance, r3 cells mix more readily with other r3 cells than with cells from other odd-numbered rhombomeres [6] – indicating that the networks controlling expression of surface receptors are not the same in all odd rhombomeres. Therefore, while it remains possible that some odd versus even features are controlled by smaller shared networks, or that a molecular periodicity becomes apparent later in development, we consider these scenarios unlikely. Instead, we favor the idea that odd versus even features are controlled by different gene networks operating in each rhombomere.

We also note that differentiating neurons cluster together (Fig. 1D, 2D), and that these clusters express low levels of AP and DV (except for *casz1*) positional genes (Fig. 1E-P; 2E, F). This suggests that neurons down-regulate positional genes when they start to differentiate. Notably, binding motifs for positional TFs (e.g., *egr2b, mafba*) remain accessible in these cells at least until 16hpf (Fig. 4C-G), perhaps indicating that changes to the chromatin state lag the transcriptional changes. Lastly, since many positional TFs maintain their expression via auto-regulatory loops, breaking such loops and down-regulating their expression to initiate differentiation may be an active process, but this remains to be tested.

### Three molecularly defined progenitor domains may correspond to pre-rhombomeres

It remains unclear how the rhombomeres arise during embryogenesis. Although the hindbrain primordium – together with the entire neural tube – becomes patterned along its AP axis via the action of various morphogens (e.g., fibroblast growth factors, retinoic acid, wnt proteins etc.; reviewed in [66]), visual observations reveal that the rhombomeres do not form in a strict AP order. In zebrafish, the r3/r4 and r4/r5 borders form first, which delineates r4 as the first rhombomere. This is followed by the r1/r2 boundary, which leaves a transient r2+r3 structure that is subsequently subdivided by formation of the r2/r3 boundary. Similarly, a transient r5+r6 segment is formed before being subdivided by the r5/r6 boundary [14, 66]. The formation of early transient segments may be evolutionarily conserved, since three potentially analogous ‘pre-rhombomeres’ have been observed in chick and four ‘primary rhombomeres’ (where the fourth corresponds to the caudal hindbrain) are present in human embryos [11–13]. Further, these early structures appear not to be solely a morphological phenomenon, but to have a genetic basis. In zebrafish *valentino* mutants (that lack *mafb* TF function), r5 and r6 do not form properly, but are replaced by an ‘rX’ rhombomere that is postulated to contain precursors for r5+r6 [14, 15]. rX has not been well defined and it is not clear if it is present in wildtype embryos, nor if there are analogous progenitors for the r2+r3 segment. Our 10hpf Multiome data identify three hindbrain populations (PHPDs), and our analyses demonstrate that they contain rhombomere progenitors.

The HB.2 population contains r5+r6 progenitors, suggesting that it corresponds to rX, while HB.1 appears to correspond to the r2+r3 segment, and HB.3 to the r4 segment – indicating that our analyses provide the first molecular characterization of the progenitor domains that precede formation of individual rhombomeres. It is likely that the PHPDs in turn arise from an earlier unpatterned primordium. Indeed, analyses by several labs have shown that blockade of all Hox TF function – by inhibiting key Hox cofactors – produces a structure that uniformly expresses early hindbrain markers [67, 68]. It remains unclear if an equivalent structure exists in wild type embryos, but scMultiome analyses at earlier stages of embryogenesis might allow a molecular characterization of such a primordium.

### PHPD cells have mixed gene expression profiles and may be multipotent

Our data reveal that the HB.1 cluster expresses markers of both r2 (*vgll3*) and r3 (*egr2b, nab1b*) at 10hpf (Fig. 3). A closer examination further demonstrated that individual HB.1 cells co-express r2 and r3 markers and such co-expression was confirmed by HCR analyses in developing embryos (Fig. 6). These findings indicate that HB.1 cells possess a mixed gene expression profile and may be able to take on both r2 and r3 fates. Accordingly, when 10hpf cells are computationally integrated with cells from 13hpf and 16hpf, they contribute to both the r2 and r3 clusters (Fig. 6). While these analyses are consistent with HB.1 cells being bipotent, this possibility awaits confirmation using lineage labeling in vivo. Nevertheless, our findings raise the question how the mixed gene expression profiles are resolved into rhombomere-specific profiles. Extensive analyses have demonstrated that cell fates along the dorsoventral axis of the neural tube are controlled by repressive interactions between TFs [69, 70]. An analogous process may be at work in the PHPDs, perhaps such that *vgll3* and *egr2b* (or components of their respective gene networks) repress each other’s transcription to force an r2 versus an r3 profile. Our HCR analyses indicate a gradual resolution of r2 versus r3 profiles along the AP axis, such that HB.1 cells at the anterior end lose *egr2b* expression first. This may indicate that a morphogen with a source in the anterior hindbrain (such as Fgf8 produced at the MHB; [71]) acts to bias *vgll3* versus *egr2b* expression within the HB.1 domain, thereby shifting the repressive balance between these TFs and resolving the mixed gene expression profile. The mixed cell identities we observe in the HB.1 domain also appear unrelated to the process of ‘border sharpening’ that takes place during rhombomere formation [72]. In that case, cells of adjacent rhombomeres intermix during boundary formation, which leads to some instances of cells in one rhombomere expressing a gene indicative of the adjacent rhombomere. Such cells are relatively few, usually associated with a rhombomere border, and become sorted into the correct rhombomere based on their expression of *Eph* and *ephrin* cell surface molecules. This contrasts with the mixed identity cells we observe in the HB.1 domain, that are more numerous and not associated with rhombomere boundaries – which have yet to form at this stage. We expect that cells in the HB.2 cluster similarly display a mixed gene expression profile between r5 and r6, but we currently lack robust r6-restricted markers to test this directly. However, we note that 10hpf HB.2 cells contribute to both r5 and r6 in the integrated UMAP (Fig. 6E) and that the rX cells found in this region of *mafB* mutant embryos give rise to both r5 and r6 [14, 15] – consistent with HB.2 corresponding to a shared progenitor pool for r5 and r6.

## ACKNOWLEDGEMENTS

This work was supported by grant NS038183 to CGS. scMultiome data has been deposited at GEO under record number GSE223535.

## AUTHOR CONTRIBUTIONS

Y-I.K. and C.S. designed all experiments. Y-I.K. carried out all bench experiments to dissect and purify hindbrain cells, to generate drug-treated samples and to assess gene expression by HCR. R.O. carried out all bioinformatics analyses in consultation with Y-I.K. and C.S. C.S. wrote the manuscript in consultation with Y-I.K. and R.O. All authors approved the final manuscript. C.S. secured funding for the project.

## METHODS

### scMultiome sample preparation

To enrich for hindbrain cells, tissue was dissected from the posterior edge of the eye to the level of somite 5 from 25 *hoxb1a:eGFP^um8^* transgenic embryos (that express GFP in rhombomere 4; [73]) at 13hpf and 16hpf. Since the hindbrain primordium is difficult to detect at 10hpf, 100 whole embryos were pooled and used at this stage. Tissue was collected in 1XPBS, dissociated by repeated pipetting through a p1000 tip, and centrifuged at 4°C at 400G for 5 minutes followed by resuspension of the pellet in 500 μl of protease solution (10 mg/ml BI protease (Sigma, P5380), 125 U/ml DNase, 2.5 mM EDTA in PBS) for 15 minutes on ice. Samples were centrifuged at 4°C at 400G for 5 minutes, resuspended in 1ml HBSS + FBS (2%), filtered through a 20 μm cell strainer (pluriSelect, KL-071912), recentrifuged at 4°C at 400G for 5 minutes, resuspended in 500 μl of HBSS + FBS (2%) and again filtered through a 20 μm cell strainer. Following centrifugation at 4°C at 400G for 5 minutes, the cells were resuspended in 200 μl PBS. For nuclei isolation, we followed the 10X Genomics recommended protocol with minor modifications. The dissociated cells were centrifuged at 900G at 4°C for 5 minutes, resuspended in 100 μl of 0.1X lysis buffer (1 mM Tris-HCl pH7.4, 1 mM NaCl, 0.3 mM MgCl2, 0.1% BSA, 0.01 % Tween-20, 0.01 % NP40, 0.001 % Digitonin (Invitrogen, BN2006), 0.1 mM DTT, 0.1 U/μl RNase inhibitor, in nuclease-free water) and incubated on ice for 5 min. Samples were then washed three times by centrifugation at 4°C at 900G for 10 min and resuspension in 1 ml of chilled wash buffer (10 mM Tris-HCl pH7.4, 10 mM NaCl, 3 mM MgCl2, 1% BSA, 0.1 % Tween-20, 1 mM DTT, 1 U/μl RNase inhibitor in nuclease-free water). Finally, nuclei were counted and resuspended in 1X nuclei buffer (provided in the 10x Genomics Single Cell Multiome ATAC Kit A; 1 mM DTT, 1 U/μl RNase inhibitor in nuclease free water) at a final concentration of approximately 2,000-3,000 nuclei/μl for sequencing on the 10X scMultiome platform.

### Single cell RNA-seq/ATAC-seq analysis

Fastq sequencing files from 10X Genomics multiomic single cell RNA-seq and ATAC-seq were processed through Cell Ranger ARC (v1.0.1) with a zebrafish GRCz11 library to obtain UMI gene expression counts and ATAC peak fragment counts. These were analyzed using the standard methods in the Signac (v1.6.0) package in R. Briefly, a Seurat Object was created from the matrix.h5 and fragments.tsv.gz files, annotated with GRCz11.v99.EnsDb, and ATAC peaks corrected by calling with macs2 (v2.2.7.1) and finally filtered to exclude cells with low total RNA expression, low total ATAC counts, high percent mitochondrial gene expression, nucleosome expression > 2 or TSS enrichment < 1. Gene expression was normalized with SCTransform and the dimensionality reduced with PCA. DNA accessibility was processed by performing latent semantic indexing. The Seurat weighted nearest neighbor method was used to compute a neighbor graph which was visualized with UMAP and clusters were annotated based on expression of marker genes. Per cell motif activities were scored with chromVar.

Significantly differentially expressed genes and differential activity motifs were identified for each cluster of cells. The Seurat Objects were integrated by finding the full intersecting ATAC peaks containing peaks in any of the three datasets and creating a new chromatin accessibility assay in each based on the full intersection peak file. The RNA-seq data was integrated with the Seurat V4 integration method and the new chromatin data integrated using Harmony followed by Seurat weighted nearest neighbor method to compute a UMAP and clusters. The cell types and UMAP embeddings for the DEAB treated 13hpf Seurat Object were predicted using the wild type 13hpf Seurat Object as a reference following the Seurat V4 reference mapping multimodal reference tutorial. The counts of cells in clusters which expressed a gene were based on cells which had a normalized expression of the gene greater than 0. Dendrograms were calculated with the Seurat BuildClusterTree function. Volcano plots were generated with EnhancedVolcano v1.12.0. scMultiome data has been deposited at GEO under record number GSE223535 and the code used to generate the data for each figure is available at github at https://github.com/rebeccaorourke-cu/Sagerstrom_zebrafish_hindbrain.

### Hybridization Chain Reaction analysis

In vivo gene expression was detected by hybridization chain reaction (HCR). HCR bundles were purchased from Molecular Instruments (https://www.molecularinstruments.com). Embryos were fixed in 4% paraformaldehyde, dehydrated with 100% methanol, and stored at −20°C until use. Embryos were rehydrated through a series of graded methanol/PBST (PBST: 1XPBS, 0.1% Tween20) washes for 5 min (75% methanol/25% PBST; 50% methanol/50% PBST; 25% methanol/75% PBST; 100% PBST). Embryos were pre-hybridized with probe hybridization buffer (provided by manufacturer) for 30 min at 37°C and then placed in hybridization buffer containing the desired HCR probe sets (2 pmol of each probe set) overnight at 37°C. The next day, the probe solution was removed, and the embryos washed with probe wash buffer (provided by manufacturer) 4 times for 15 min at 37°C, followed by washes in 5X SSCT (5X SSC, 0.1%Tween20) 2 times for 5 min. Embryos were pre-treated in amplification buffer for 30 min at room temperature. Separately, hairpin h1 and h2 were prepared by snap cooling a 3 uM stock, heating at 95°C for 90 seconds and cooling to room temperature in a dark drawer for 30 min. h1, h2 hairpins were then added to amplification buffer and incubated with embryos overnight in the dark at room temperature. The next day, the hairpin solution was removed, embryos were washed with 5X SSCT and kept at 4°C until imaging.

### DEAB and SU5402 treatments

4-Diethylaminobenzaldehyde (DEAB) was used to inhibit retinoic acid signaling. DEAB (Sigma, D86256) was dissolved in DMSO and 5 μM DEAB was used to treat embryos from 4hpf until collected for analysis. SU5402 (Abcam, ab141368) was used to inhibit FGF signaling. SU5402 was dissolved in DMSO and 10 μM used to treat embryos from 4hpf until collected for analysis.

---

**Figure S1.**
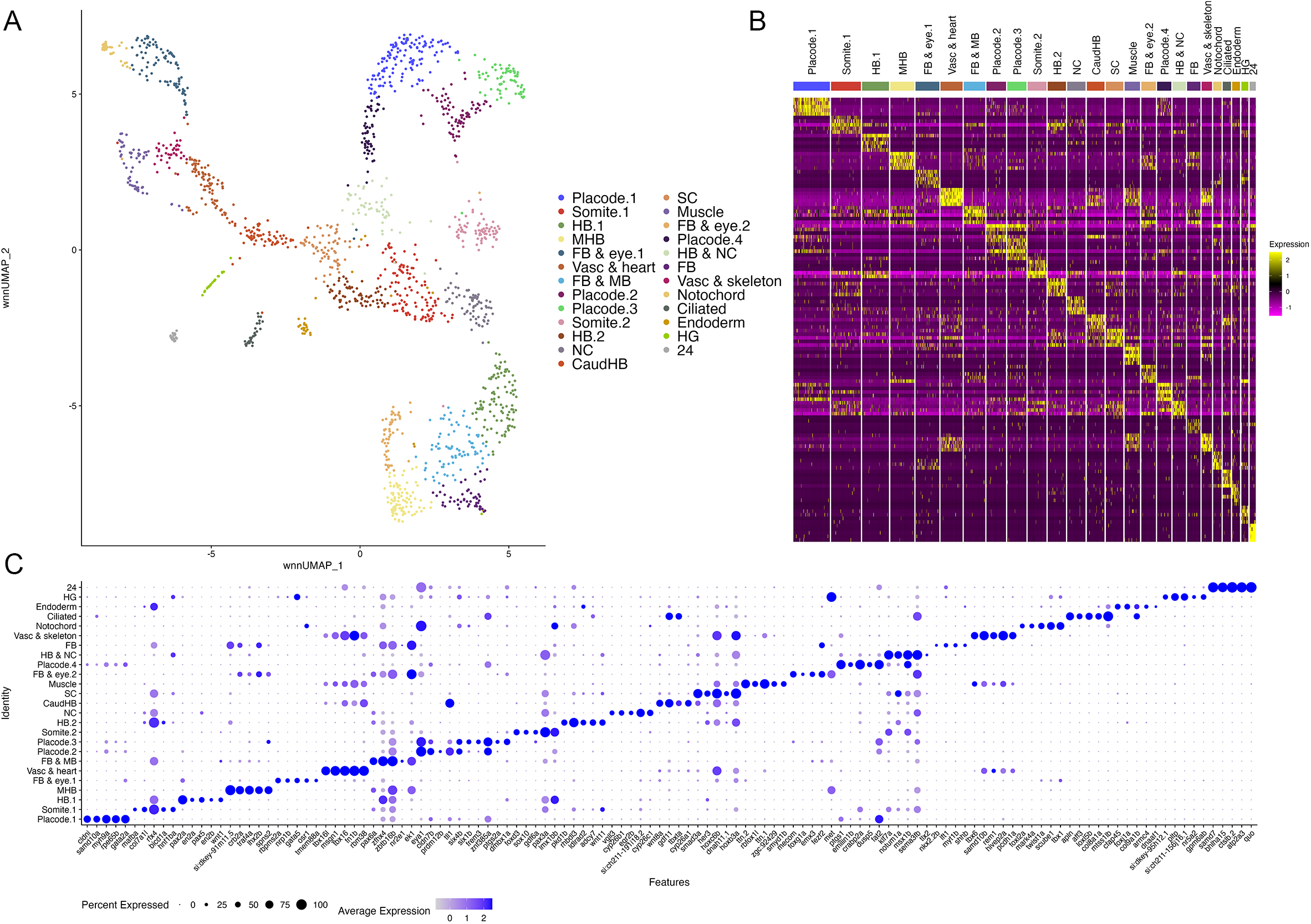
scMultiome analysis of 10hpf zebrafish. A. UMAP showing all clusters obtained from 10hpf whole zebrafish embryos. B. Heatmap showing expression of the top 5 enriched genes in each cluster. C. Dot plot showing expression ofthe top 5 enriched genes in each cluster.

**Figure S2.**
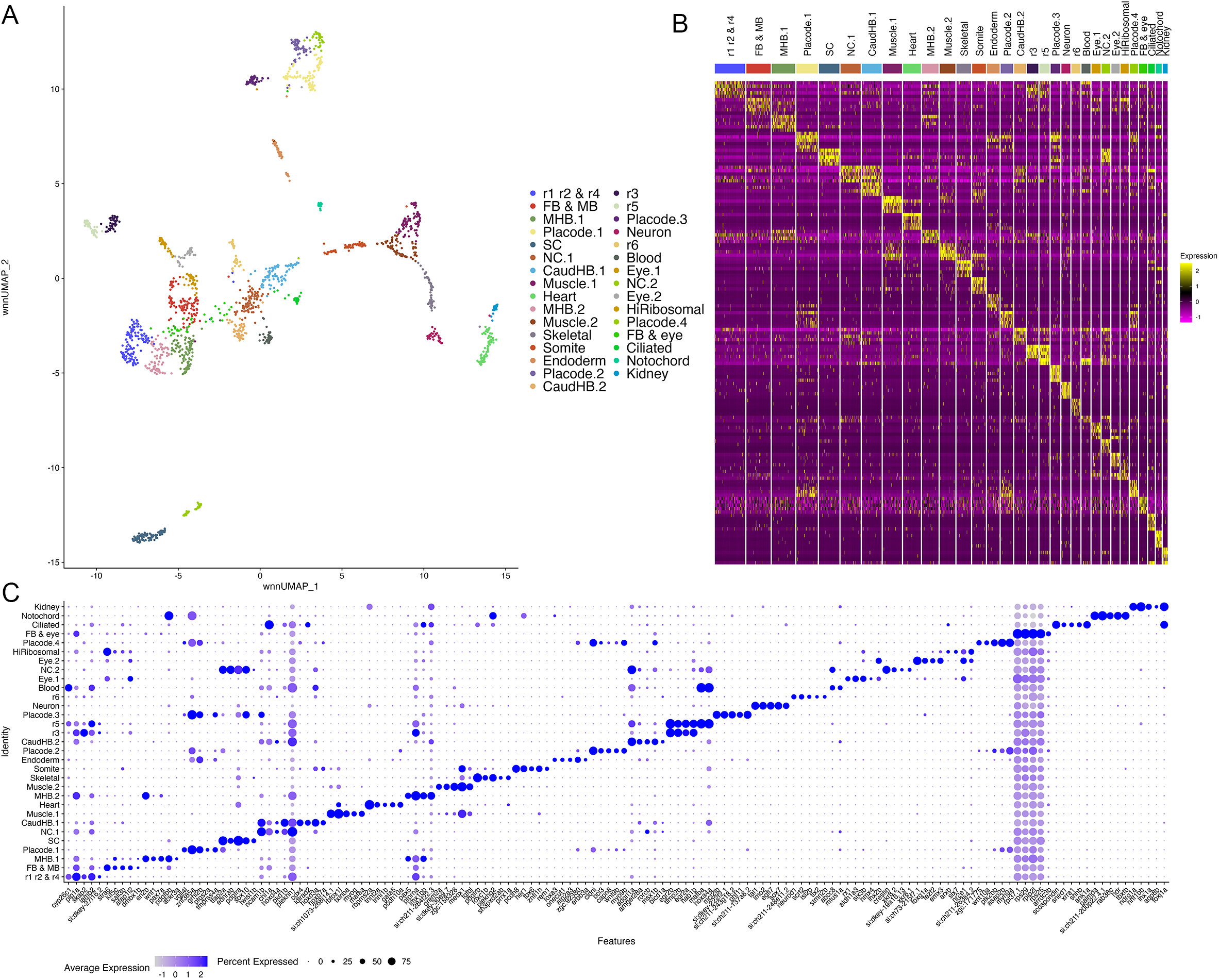
scMultiome analysis of 13hpf zebrafish. A. UMAP showing all clusters obtained from 13hpf dissected zebrafish hindbrain regions. B. Heatmap showing expression of the top 5 enriched genes in each cluster. C. Dot plot showing expression of the top 5 enriched genes in each cluster.

**Figure S3.**
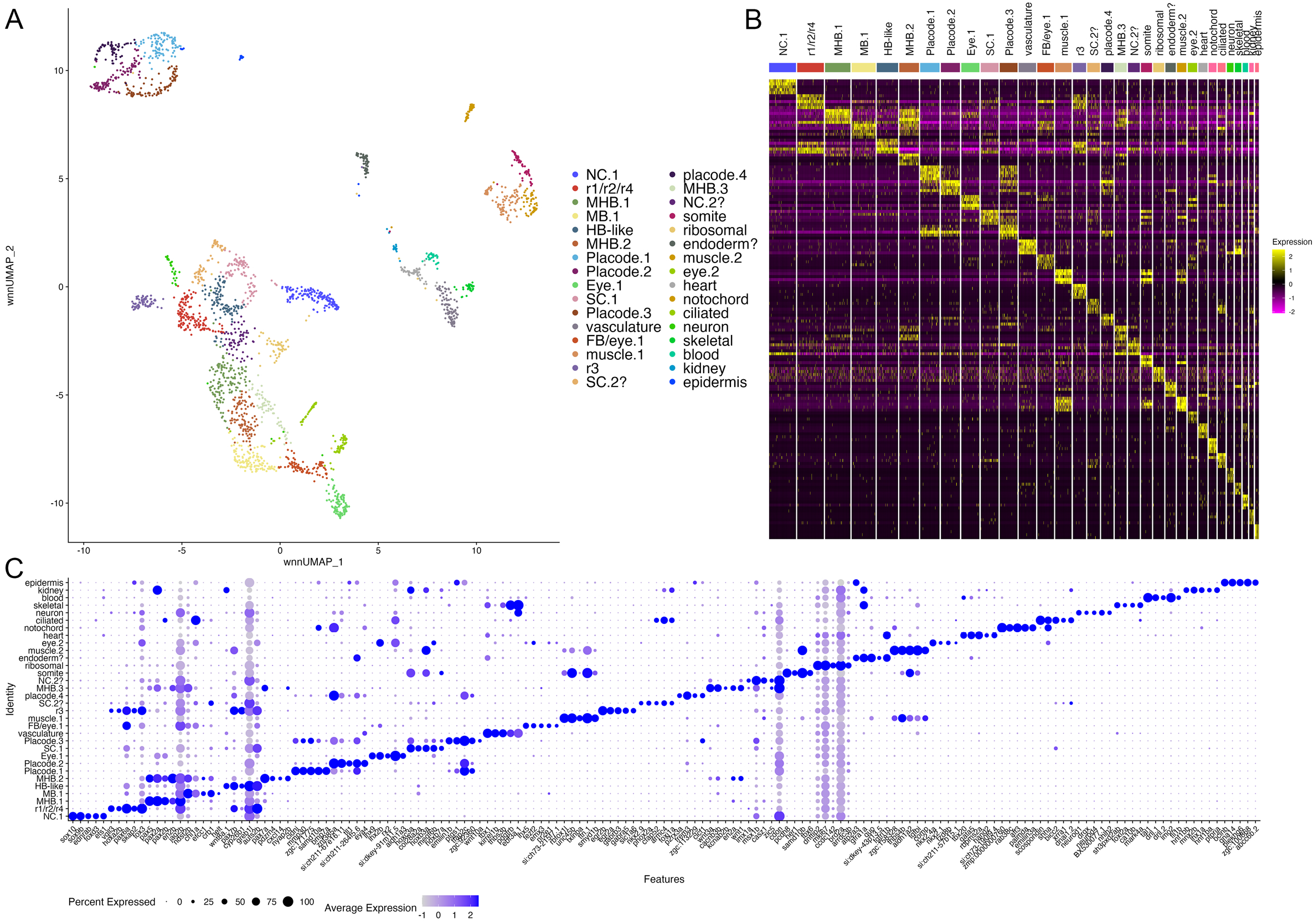
scMultiome analysis of DEAB-treated 13hpf zebrafish. A. UMAP showing all clusters obtained from 13hpf dissected zebrafish hindbrain regions treated with DEAB. B. Heatmap showing expression of the top 5 enriched genes in each cluster. C. Dot plot showing expression of the top 5 enriched genes in each cluster.

**Figure S4.**
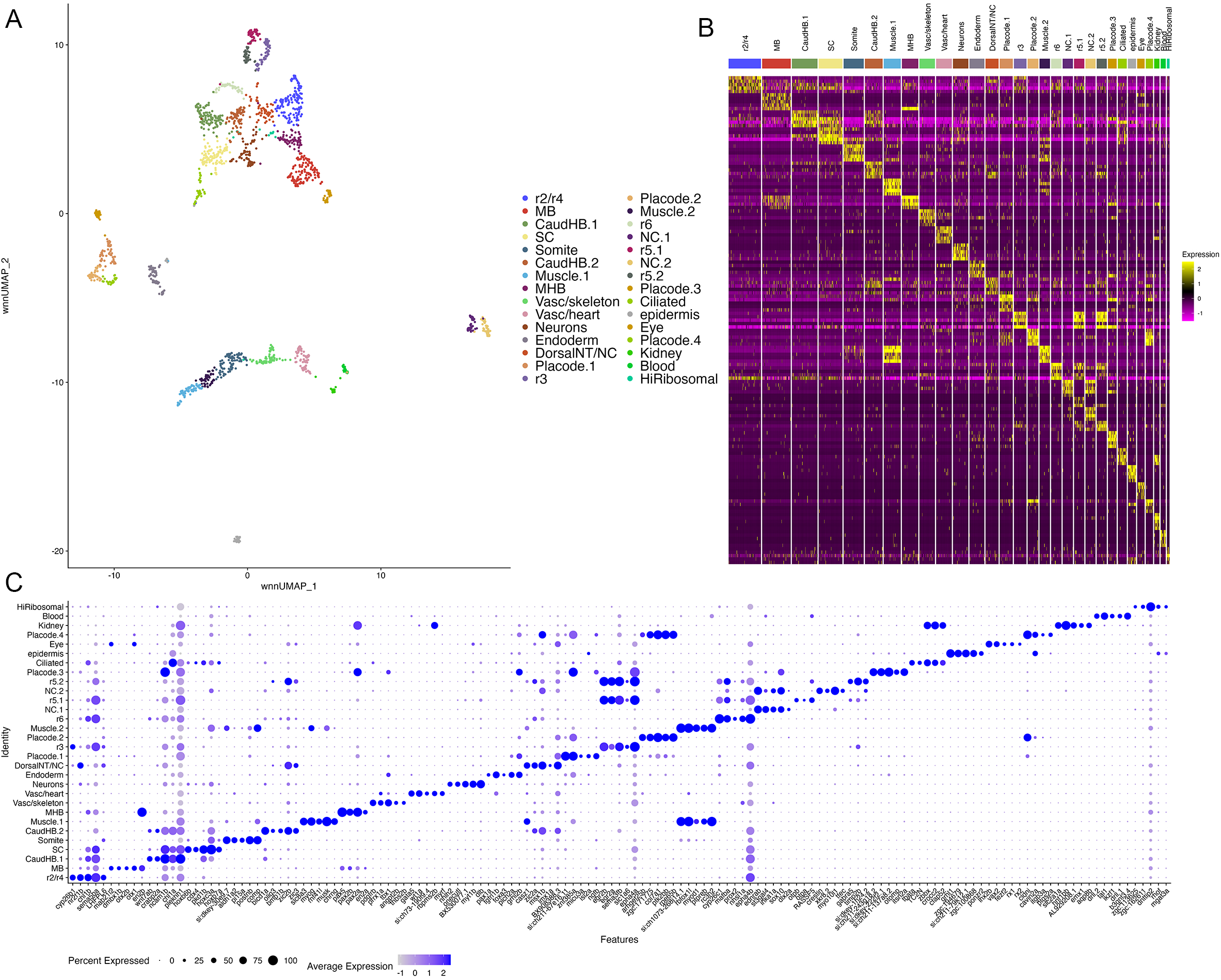
scMultiome analysis of 16hpf zebrafish. A. UMAP showing all clusters obtained from 16hpf dissected zebrafish hindbrain regions. B. Heatmap showing expression of the top 5 enriched genes in each cluster. C. Dot plot showing expression of the top 5 enriched genes in each cluster.

**Figure S5.**
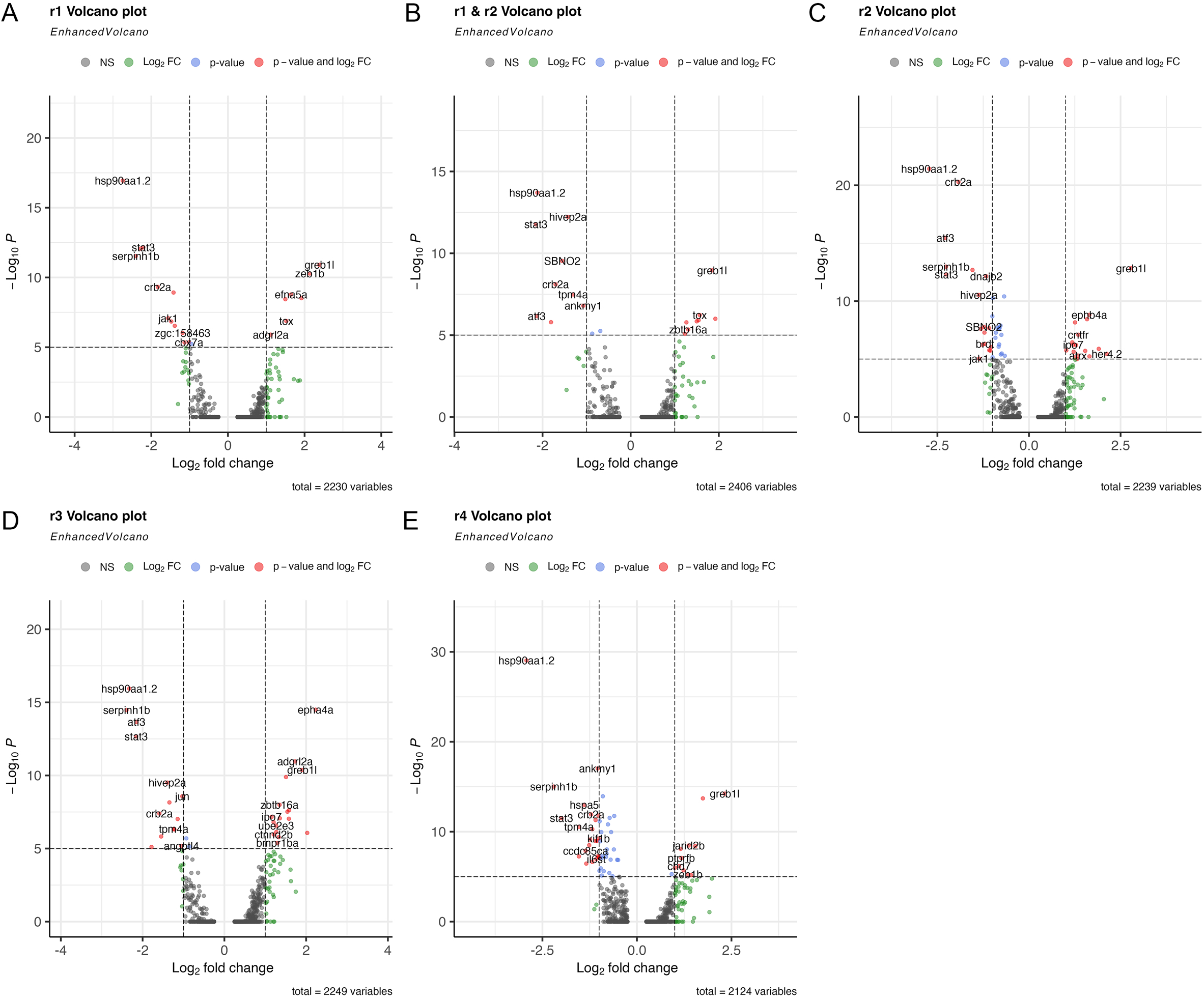
DEAB-treatment has a limited effect on gene expression in rhombomeres 1-4. A-E. Volcano plots showing differential gene expression between control and DEAB­ treated cells for each rhombomere. A positive log2FC indicates higher expression in control than DEAB.

